# Molecular mechanisms of heart field specific cardiomyocyte differentiation- a computational modeling approach

**DOI:** 10.1101/2024.12.19.629328

**Authors:** Georgios Argyris, Ricco Zeegelaar, Janine N. Post

## Abstract

Tissue engineering protocols achieve building miniature hearts but mechanisms determining cell differentiation still need to be fully understood and optimized. In this study, we present a gene regulatory network (GRN) that describes the differentiation of committed *cardiomyocytes* towards ventricular or atrial cardiomyocytes. The GRN is coupled with Boolean dynamics and steady state analysis shows steady states which agree with the experimental expression of marker genes. Our Boolean model extends earlier work on a model describing the first and second heart field formation to include atrial and ventricular cardiomyocytes. Thus, our study paves the way for the generation of heart field-specific cardiomyocytes located in specific chambers of the fully developed heart. The Boolean model is validated through simulations and by its ability to reproduce known knockouts.

## 1 Introduction

The heart consists of multiple cell types, including endothelial cells, fibroblasts, and cardiomyocytes. To achieve effective tissue replacements in cardiac defects, it is critical to direct differentiation toward specific cells with the same properties.. Even between cardiomyocytes (CM), gene expression varies depending on their location within the heart, and researchers strive to produce these subtypes of CM in vitro. Consequently, there is an urgent need to study the mechanisms that determine cell differentiation. Cell differentiation is governed by gene regulatory networks (GRNs) whose dynamics can be explored by Boolean logic models [1, 11].

In this study, we present a GRN coupled with Boolean dynamics of a coarse cardiomyocyte developmental lineage. The lineage includes two developmental stages: the *heart field formation* (*“Heart Field Identity”* box of Fig. 1) and the *CM differentiation* (*“Cell Identity”* box of Fig. 1). Heart field formation occurs at the start of the developmental process. After mesoderm formation, cardiogenic mesodermal progenitors migrate and form two heart fields [5]: the *first heart field (FHF)*, and the *second heart field (SHF)*. In [11], Herrmann et al. introduce a Boolean Network (BN) which describes the early murine cardiac development. Their BN exhibits two steady states with different gene expression where each steady state corresponds to a specific heart field (see *Heart Field Identity* box of Fig. 1). Both heart fields contribute to the formation of CMs that populate the muscular wall of the heart chambers: the *Atrium* and the *Ventricle* [11, 17]. Specifically, cells of the first heart field primarily populate the left ventricle, cells of the second heart field populate the right ventricle, and both fields contribute to the atria [17] (see *Cell Identity* box of Fig. 1).

**Fig. 1:**
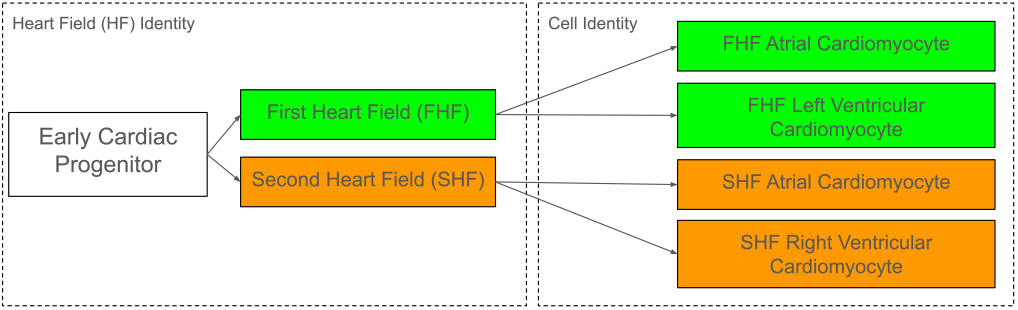
The coarse cardiac developmental lineage. (Left Box) Cardiac progenitor cells form the two heart fields. (Right Box) The heart fields contribute to the formation of cardiomyocytes. A more detailed Figure can be found in Appendix A.

Atrial and ventricular CMs are functionally distinct and are characterized, among others, by expression of different transcription factors [26]. The mechanisms behind the differential expression between the types of CMs are still not well understood, limiting the capabilities for directed in vitro differentiation. To help experimentalists towards this goal, *(i)* we derive a prior knowledge network of the GRN representing *CM differentiation, (ii)*we then couple the GRN with Boolean dynamics and, finally *(iii)* we validate the model through attractor analysis. The *cardiomyocyte BN* reproduces exactly two steady states: one corresponding to the atrial CM and one corresponding to the ventricular CM.

Due to differences in gene expression profile and differentiation potential [11, 43], it is beneficial to guide cells not only to a specific atrial or ventricular cell fate, but also through a specific heart field. To address this challenge, we merge the Heart Field BN of [11] with the cardiomyocyte BN. The merged BN exhibits six steady states in general, four of them being biologically relevant since they correspond to distinct cell types (see *Cell Identity* box in Fig. 1). We perform simulations of a probabilistic version of the BN [33] to ensure that most trajectories reach biologically plausible steady states (distinct cell types). The simulation analysis of the merged BN sketches the framework for generation of heart-field-specific atrial and ventricular CMs. The merged BN also reproduces several in silico knockout and overexpression experiments.

Our contributions are threefold and constitute the three sections of the manuscript: in Section 2 we present Boolean network of cardiomyocyte differentiation, in Section 3 we merge the CM BN with an existing BN that models heart field formation, and in Section 4 we analyze the merged BN with simulations and in silico gene perturbation experiments.^1^

## 2 A Boolean Network of the Cardiomyocyte

In this Section we introduce a Boolean Network (BN) of the cardiomyocyte. Particularly, in Section 2.1 we provide and describe a GRN, and in Section 2.2 we couple it with Boolean dynamics.

### 2.1 A Gene Regulatory Network of the Cardiomyocyte

The first information that modellers obtain about a GRN is usually encoded into an Interaction Graph (IG). The IG consists of nodes and edges where nodes represent genes, proteins, chemical compounds and so on, operating inside the cell. The edges represent the effects between the nodes such as activation, inhibition and repression. Such an IG can be constructed either by extensive literature study (called *prior knowledge network* (PKN)) or can be derived from data. Here, we construct a PKN by selecting literature on general CM gene expression, differential structural gene expression, and differential transcription factor expression. The resulting PKN is presented in the left part of Fig. 2.

**Fig. 2:**
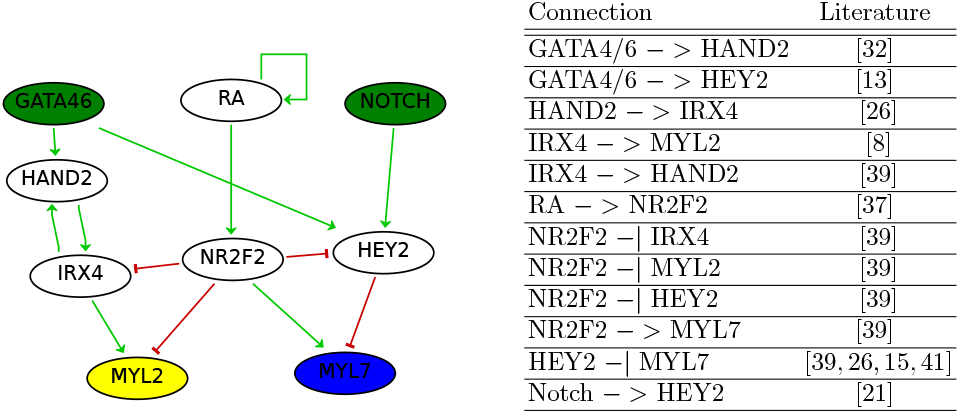
(Left) An IG representing the cardiomyocyte (drawn in GinSim [23]). The blue node *MYL7* is an atrial marker and the yellow node *MYL2* is an ventricular marker. The green nodes *GATA4/6, NOTCH* are both active since a progenitor cell state. (Right) Table with the relevant literature of the influences. An extended version of this table can be found in Appendix B.

The PKN of Fig. 2 constitutes a static representation [7] of the differentiation process that describes the interactions between genes (i.e., positive or negative influence) when a cell commits to either a ventricular or atrial CM. In particular, the graph proposes how the signaling molecule retinoic acid (node *RA*) works. is combined with the regulatory network to affect the expression of transcription factors that regulate the distinction between atrial and ventricular CMs. The *NR2F2* gene, which encodes the *COUPTF-II* protein, possesses a central role in the GRN. RA upregulates *NR2F2* whose activity eventually orchestrates the decision making process.

On top of Fig. 2, the green nodes constitute genes which are assumed to always be expressed in premature CMs. Although *GATA4* and *GATA6* have slightly different roles in cardiac development, their role in the CM can be summarized in one single node, like in [11]. The node *GATA4/6* is considered to be always active because a threshold of *GATA4* and *GATA6* expression is required for cardiovascular development according to [40]. Indeed, *GATA4/6* exhibits activity in both the first and second heart field during murine cardiac development [11]. It has been shown that the interaction between *GATA* transcription factors, *NOTCH*, and the absence of *NR2F2* constitutes the mechanism underlying the expression of *HEY2* [13]. Therefore, we conclude that *NOTCH* signaling has to be active.

At the bottom of the Fig. 2, the yellow node (gene *MYL2*) is a marker of a ventricular CM while the blue node (gene *MYL7*) is a marker of an atrial CM. The retinoic acid (node *RA* on top of the figure) is the driver of the system; when *RA* is activated, *MYL2* is inhibited while *MYL7* is activated leading the CM to an atrial identity. Although we use *MYL7* as an atrial marker, it would be plausible to use other markers like *SLN* or *MYL4* which are expressed only in atria [26].

Note that certain interactions in the GRN of Fig. 2 are not direct, but are a result of simplifications. To assess the reliability of the GRN, we have to incorporate a compatible dynamical model capable of reproducing the experimental observations. If the GRN is accurate enough, a dynamical model will be able to unravel the empirical evidence of cell differentiation: the transition from a committed CM towards atrial and ventricular CMs.

### 2.2 Boolean Modelling of the Cardiomyocyte GRN

To investigate the accuracy of the GRN (left part of Fig. 2), we equip it with Boolean dynamics. BNs are established dynamical formalisms that receive much attention due to their simplicity. In a BN, the variables take binary values, i.e. *0,1* (or *False, True*) that denote whether the corresponding node is active or not. We propose that the BN on the left part of Fig. 3 properly models the differentiation procedure of a committed CM to either ventricular or atrial CM. Before proceeding with the validation of the BN, we have to introduce some preliminary notions.

**Fig. 3:**
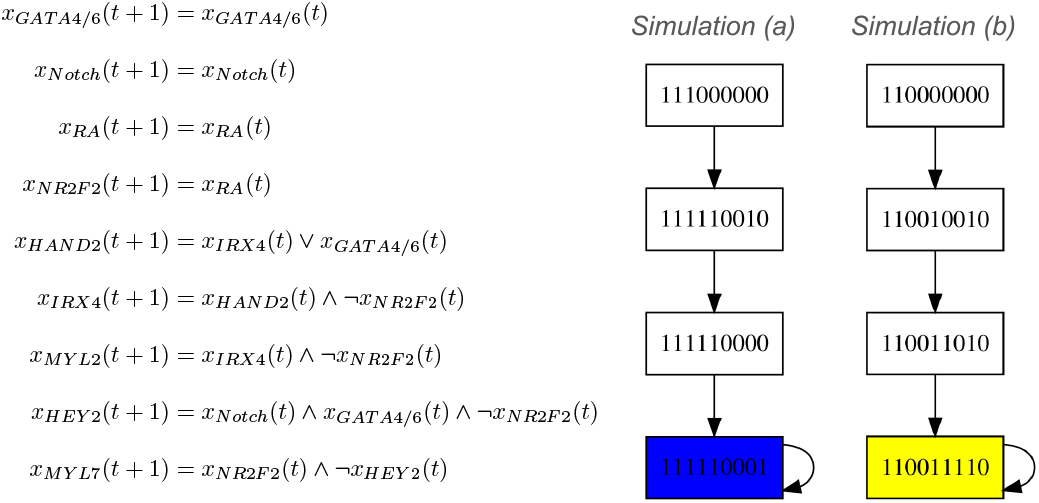
(Left) A BN compatible with the IG of Fig. 2. (Right) Simulations.

#### Transition

At a specific time point *t*, let the variables (which represent the activity of the GRN nodes) have the following values:

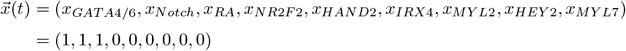

A vector of such values is called *state*. By evaluating the BN of Fig. 3, we obtain the vector 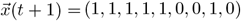. Therefore, we say that there exists a transition from one state to another and we denote it with 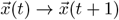.

#### Simulation and State Transition Graph

We may have whole sequences of transitions by applying the update functions iteratively. These sequences of states and transitions are called *simulations* (in literature, the notion of simulation can be found with several different names such as solutions, orbits, and trajectories, especially in the case of continuous time dynamical systems). We present two simulations starting from different initial states at the right part of Fig. 3. All simulations are encoded into the State Transition Graph (STG) which contains all states and all transitions between the states. However, if a BN consists of *n* variables, the STG contains *2* ^*n*^ states and, therefore, it is impossible to display it for large values of *n*.

#### Update modes

It is worth mentioning that, in the simulations of Fig. 3, the updates of all variables are performed synchronously and, therefore, each state (box of Fig. 3) has exactly one successor state. Different update modes reproduce different transition sets. For the rest of the article, we consider the asynchronous update mode wherein only one variable is non-deterministically (randomly) selected and updated (see example in Fig. 4). This is considered as one of the most biologically plausible update modes because of the different timescales involved in the various biological processes. For example, changing gene expression programs can take hours or even days, protein complex formation goes on the second scale, and post-transcriptional protein modifications take minutes to happen [30].

**Fig. 4:**
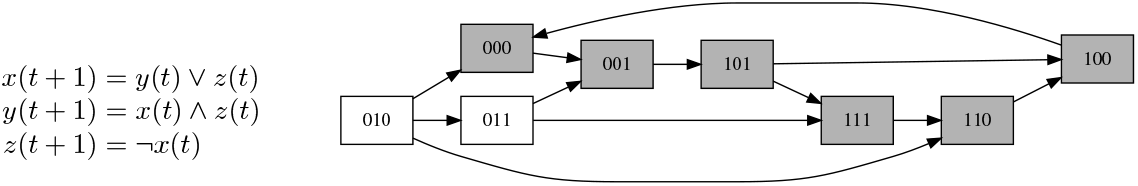
(Left) A BN. (Right) The STG generated by the asynchronous update mode wherein only one variable is non-deterministically selected and updated. Self-transitions of all states are not displayed.

#### Attractors

We start by noticing that, when the initial condition of the variable RA is *1* (*simulation (a)* of Fig. 3), the BN evolves and remains at a state (the blue state) where *MYL7* is *on* (*1*) and *MYL2* is *off* (*0*). Hence, the cell becomes an atrial CM, as observed in vitro [31, 39]. In contrast, if the simulation starts from a state where RA is off (*simulation (b)* of Fig. 3), the BN evolves and remains at a state (the yellow state) where MYL2 is *on* and MYL7 is *off*. Thus, the cell becomes a ventricular CM. The yellow and blue states are called steady states and, traditionally, in cell differentiation processes different steady states correspond to different cell types. A BN may also exhibit cyclic attractors; attractors with more than one state where the BN evolves and oscillates between. In the right part of Fig. 4, we display a cyclic attractor which consists of the grey sequences of states.

#### Validation

We use the tool Boolsim [9], incorporated in the CoLoMoTo Notebook [25], to compute the attractors of the asynchronous BN. We stabilize the variables *x*_*GATA4/6*_ and *x*_*Notch*_ by setting their functions equal to the value *1* because these variables are already active in the progenitors, and stay active until the end of the developmental process. The Table 1 displays the two attractors of the BN, both of them being steady states.

**Table 1:** The two steady states of the BN of Fig. 3.

|  | GATA4/6 | Notch | RA | NR2F2 | HAND2 | IRX4 | MYL2 | HEY2 | MYL7 |
| --- | --- | --- | --- | --- | --- | --- | --- | --- | --- |
| aCM type | 1 | 1 | 1 | 1 | 1 | 0 | 0 | 0 | 1 |
| vCM type | 1 | 1 | 0 | 0 | 1 | 1 | 1 | 1 | 0 |

Specifically, the rows display the two steady states. In each steady state, we see if the corresponding node is active or inactive: if an entry contains *1* (green), the node of this column is active in the steady state while, if an entry contains *0* (cyan), the relevant node is inactive.

We see that the results are compatible with the known marker genes for ventricular and atrial CMs [31, 39]: the BN exhibits exactly two steady states, one corresponding to the ventricular CM where the relevant nodes (*GATA4/6, Notch, HAND2, IRX4, MYL2, HEY2*) are active, while the second steady state corresponds to the atrial CM where the relevant atrial nodes are active (*GATA4/6, Notch, HAND2, NR2F2, MYL7*). Depending on the initial condition of the BN and, in particular on the initial condition of *RA*, the cell becomes either an atrial or a ventricular CM.^2^

## 3 Merging the Cardiomyocyte BN with a BN that Determines First and Second Heart Field Identity

To cover the entire lineage from early cardiac progenitors to CMs, we merge the cardiomyocyte BN (of Section 2) with the only BN of the literature that describes heart development. More specifically, in Section 3.1 we present and study (under the asynchronous update mode) the BN of [11] that describes heart field formation. In Section 3.2 we merge the heart field BN of Section 3.1 with the cardiomyocyte BN and perform attractor analysis.

### 3.1 A BN that Determines First and Second Heart Field Identity

In [11], Herrmann et al. presented a BN model that includes interactions between genes during the early murine cardiac development. They focused on two areas with different gene expression, namely, the first heart field (FHF) and the second heart field (SHF). Two steady states of their BN correspond to the two different heart fields which are characterized by the expression of specific genes. We present the model in Fig. 5. The node *exogen BMP2* (*ex_BMP2*) is green because its initial condition is always *1* since the differentiation of CMs begins at lateral plate mesoderm where *ex BMP2* is always active [29].

**Fig. 5:**
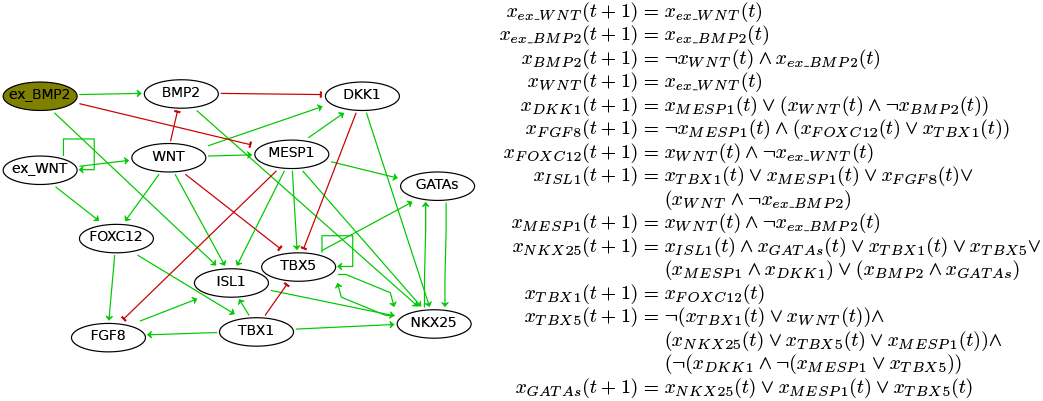
(Left) A GRN that describes the differentiation between the FHF and the SHF taken from [11]. (Right) The BN taken from [11].

#### Configuration and Hypothesis

We repeat the attractor analysis of [11] where the authors compute the steady states through exhaustive simulations using synchronous updating (as described in the simulations of the right part of Fig. 3). On the contrary, we motivate asynchronous updating where only one node is non-deterministically selected and updated at a time step. To investigate the fate of attractors in the asynchronous update mode, we utilize the tool Boolsim [9]. Our crucial hypothesis is that the attractors are exactly the same, independently of the updating dynamics.

#### Attractor Analysis

We present the *3* attractors, all of them being steady states, in Table 2. The first column of Table 2 contains the names of the steady states while the first row contains the name of the nodes. If an entry contains *1* (green), the node of the corresponding column is active in the steady state of the row, while if the entry contains *0* (cyan) the relevant node is inactive in this steady state.

**Table 2:** The three attractors of the BN of Fig. 5.

|  | ex_WNT | ex_BMP2 | BMP2 | WNT | DKK1 | FGF8 | FOXC12 | ISL1 | MESP1 | NKX25 | TBX1 | TBX5 | GATAs |
| --- | --- | --- | --- | --- | --- | --- | --- | --- | --- | --- | --- | --- | --- |
| FHF | 0 | 1 | 1 | 0 | 0 | 0 | 0 | 0 | 0 | 1 | 0 | 1 | 1 |
| SHF | 1 | 1 | 0 | 1 | 0 | 1 | 1 | 1 | 0 | 1 | 1 | 0 | 1 |
| null | 0 | 1 | 1 | 0 | 0 | 0 | 0 | 0 | 0 | 0 | 0 | 0 | 0 |

#### Interpretation

The two different steady states correspond to the first heart field (FHF) and the second heart field (SHF). Remarkably, the steady states remain unchanged regardless of the update mode while the asynchronous update mode does not generate any additional attractors. Briefly, our attractor analysis and the attractor analysis of [11] are producing the same attractor set. Indeed, the latter was expected because: *(i)* if a state is steady under the synchronous update mode, it is also steady under any other update mode, and *(ii)* most cyclic attractors in BNs are artifacts of the synchronous update mode and disappear in the case of asynchronous dynamics [4].

### 3.2 Merging the Heart Field with the Cardiomyocyte BN

In this section, we merge the heart field BN and the cardiomyocyte BN. Our crucial hypothesis is that the coupling of the two BNs and the exploration of attractor fate in the merged BN will provide interesting insights in the underlying mechanism of generating heart field specific atrial and ventricular CMs.

#### Configuration

The only common node between the CM GRN and the heart field GRN is the node that represent the *GATA* genes. These genes are expressed in heart development a priori. In addition, the *GATA* node is active in all steady states of both the heart field BN and the cardiomyocyte BN (we correctly considered that the *GATA4/6* node is always active). Therefore, we connect the heart field BN and the cardiomyocyte BN up to the *GATAs* node and the *GATA4/6* node respectively. We present the merged BN in Fig. 6.

**Fig. 6:**
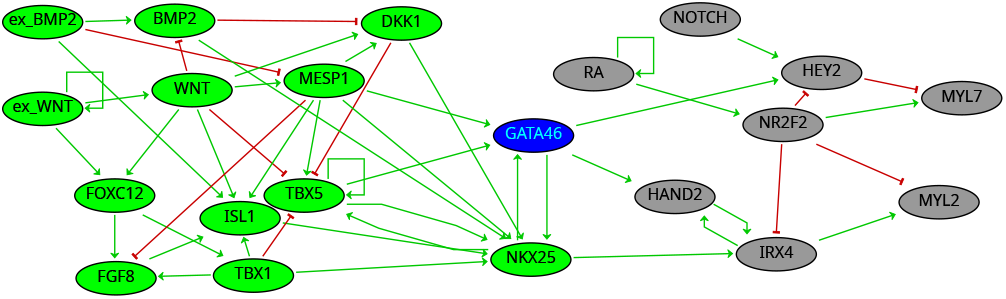
The merged GRN. The nodes in lime color (mostly left part of Fig. 6) constitutes the heart field GRN taken from [11] while in grey colour are the nodes of the Cardiomyocyte GRN. The node in blue colour (*GATA4/6*) is the (overlapping) connective node and represents the *GATA* genes.

The unification of the two BNs is performed according to two considerations. Firstly, in the merged model, the update function of the *GATA4/6* node (blue node of Fig. 6) equals the *GATAs* node of the heart field BN model. In other words, in contrast with the cardiomyocyte BN where *x*_*GATA4/6*_ (*t* + *1*) = *x*_*GATA4/6*_ (*t*), the update function of *x*_*GATA4/6*_ in the merged BN (blue node of Fig. 6) is:

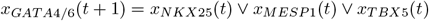

Secondly, we also incorporate one more essential activating arrow from *NKX2*.*5* towards *IRX4* (*NKX2*.*5* − > *IRX4*). *NKX2*.*5* constitutes a general cardiac gene highly expressed in both atrial and ventricular CMs, and affects *IRX4* [26]. The function of *IRX4* is the following:

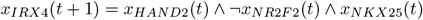

The update functions of all other variables remain the same as in the single BNs. We provide the update functions of the full BN in Appendix D.

#### Attractor Analysis

We use Boolsim [9] to compute the attractors of the merged BN under the asynchronous schema. We provide the 6 attractors, all of them being steady states, in Table 3. The first column displays the name of the nodes while the row displays the names of the steady states. If an entry contains *1*, the node of the row is active in the steady state of the column while if a table entry contains *0*, the node of the row is inactive in the steady state of the column.

**Table 3:** The 6 attractors of the merged BN of Fig. 6.

|  | FHF aCM | FHF vCM | SHF aCM | SHF vCM | null 1 | null 2 |
| --- | --- | --- | --- | --- | --- | --- |
| ex_WNT | 0 | 0 | 1 | 1 | 0 | 0 |
| ex_BMP2 | 1 | 1 | 1 | 1 | 1 | 1 |
| BMP2 | 1 | 1 | 0 | 0 | 1 | 1 |
| WNT | 0 | 0 | 1 | 1 | 0 | 0 |
| DKK1 | 0 | 0 | 0 | 0 | 0 | 0 |
| FGF8 | 0 | 0 | 1 | 1 | 0 | 0 |
| FOXC12 | 0 | 0 | 1 | 1 | 0 | 0 |
| ISL1 | 0 | 0 | 1 | 1 | 0 | 0 |
| MESP1 | 0 | 0 | 0 | 0 | 0 | 0 |
| NKX25 | 1 | 1 | 1 | 1 | 0 | 0 |
| TBX1 | 0 | 0 | 1 | 1 | 0 | 0 |
| TBX5 | 1 | 1 | 0 | 0 | 0 | 0 |
| GATA4/6 | 1 | 1 | 1 | 1 | 0 | 0 |
| Notch | 1 | 1 | 1 | 1 | 1 | 1 |
| RA | 1 | 0 | 1 | 0 | 0 | 1 |
| NR2F2 | 1 | 0 | 1 | 0 | 0 | 1 |
| HAND2 | 1 | 1 | 1 | 1 | 0 | 0 |
| IRX4 | 0 | 1 | 0 | 1 | 0 | 0 |
| MYL2 | 0 | 1 | 0 | 1 | 0 | 0 |
| HEY2 | 0 | 1 | 0 | 1 | 0 | 0 |
| MYL7 | 1 | 0 | 1 | 0 | 0 | 1 |

#### Interpretation

Since the heart field BN exhibits *3* steady states and the cardiomyocyte BN exhibits *2* steady states, the merged BN exhibits *3 × 2* = *6* steady states. In the table of Fig. 3, we notice that: *(i)* the *2nd* column (*FHF aCM*) displays a steady state that corresponds to an atrial CM derived from the FHF, *(ii)* the *3rd* column (*FHF vCM*) displays a steady state which corresponds to a ventricular CM derived from the FHF, *(iii)* the *4th* column (*SHF aCM*) displays a steady state which corresponds to an atrial CM derived from the SHF, and *(iv)* the *5th* column (*SHF vCM*) displays a steady state which corresponds to a ventricular CM derived from the SHF. The two *null* steady states (last two columns of the table) are consequences of the null steady state of the heart field BN. Indeed, in these two steady states the *GATA4/6* node, whose activity is a prerequisite for the development of heart [40], is inactive. In Section 4.2, we conduct simulations and show that, mostly, the simulations end up in the biologically relevant steady states.

The steady states in the first 4 columns of Fig. 3 (*FHF aCM, FHF vCM, SHF aCM, SHF vCM*) are in agreement with biological evidence; both types of CMs can be derived from either heart field [11, 17]. Particularly, cells of the first heart field primarily populate the left ventricle, cells of the second heart field populate the right ventricle, and both fields contribute to the atria [17]. Our analysis suggests that the steady state of the *3rd* column (*FHF vCM*) represents a left-ventricular CM while the steady state of the *5th* column (*SHF vCM*) represents a right-ventricular CM.

In Fig. 3, we observe the activity of signalling pathways (*WNT, BMP, RA, FGF* and *NOTCH*) in each steady state. The consideration of pathways’ activity is crucial for guiding the differentiation of precursor CMs towards heart-field-specific CMs when, for instance, we need to reconstruct tissue for a degenerative disease affecting the right ventricle. Our findings suggest that SHF-derived right ventricular CMs can be generated from precursor CMs, by activating the *WNT, BMP, FGF* and *NOTCH* pathway while inhibiting *RA* signalling.

We note that heart field decision takes place before the CM type decision. Thus, we cannot guarantee that the activity of the heart field nodes remain the same after the CM decision. CM nodes, which become active at a later stage, may switch off (on) several heart field nodes. In fact, steady states mostly reflect which pathways and transcription factors have had to be activated or inhibited at a specific differentiation step of the lineage to arrive at the final cell type of the model. The merging of the GRNs (and the BNs) is imperfect, and several connections are missing. Still, the merged BN can reproduce several well-known knockout and overexpression experiments that we perform in Section 4.1.

#### Formal Interpretation of Steady State Configuration

Let **𝒜**_*i*_, **𝒜**_*j*_ be two steady states, and let **𝒜**_*i,j*_ = **𝒜**_*i*_ |**𝒜**_*j*_ be the concatenation of the two steady states up to the last digit of **𝒜**_*i*_ and the first digit of **𝒜**_*j*_. We explain this notation in the following example:

#### Example 1

Consider the FHF steady state **𝒜**_*FHF*_ = (*0, 1, 1, 0, 0, 0, 0, 0, 0, 1, 0, 1*, **1**) (first row of Table 2) and the atrial steady state **𝒜**_*aCM*_ = (**1**, *1, 1, 1, 1, 0, 0, 0, 1*) (first row of Table 1). The concatenation of the steady states is the following:

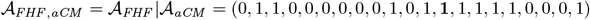

The bold digit is the steady state of the *GATAs* node in the heart field BN and the steady state of the *GATA4/6* node in the cardiomyocyte BN.

The merged BN reproduces the steady states of the single BNs in the following way. Let **𝒜**_*FHF*_ be the steady state of the FHF (first row of Table 2), **𝒜**_*SHF*_ be the steady state of the SHF (second row of Table 2), **𝒜**_*null*_ be the null steady state (third row of Table 2), **𝒜**_*aCM*_ be the the steady state of the atrial CM (first row of Table 1), and **𝒜**_*vCM*_ be the steady state of the ventricular CM (second row of Table 1). The 4 important steady states can be derived by the following concatenation rules: **𝒜**_*FHF,aCM*_ = **𝒜**_*FHF*_ |**𝒜**_*aCM*_, **𝒜**_*FHF,vCM*_ = **𝒜**_*FHF*_ |**𝒜**_*vCM*_, **𝒜**_*SHF,aCM*_ = **𝒜**_*SHF*_ |**𝒜**_*aCM*_, **𝒜**_*SHF,vCM*_ = **𝒜**_*SHF*_ |**𝒜**_*vCM*_. The other two steady states do not follow this rule as they originate from a stable condition wherein the node *GATA4/6* remains inactive (*x*_*GATA4/6*_ = *0*), while, for the cardiomyocyte BN, we consistently assume that *x*_*GATA4/6*_ = *1*.

## 4 Analysis of the Merged Boolean Network

To analyze the model’s ability to recapitulate known perturbations (knockout and overexpression experiments), we analyze the merged BN. To investigate the ability of the BN to transition random initial conditions to biologically plausible steady states, we conduct probabilistic simulations for populations of cells.

### 4.1 The Merged BN Reproduces Known Perturbations

Here, we further validate our model in terms of its ability to reproduce well-known knockout and overexpression experiments found primarily in [26, 34]. The knockout of a certain gene, also represented by deactivation or removal of the protein, is modelled in the BN by setting the value of that node to zero. For example, to investigate the knockout of the *HAND2* gene, we fix its value in the merged BN with *0* (i.e. *x*_*HAND2*_ (*t* + *1*) = *0*) while keeping all other nodes unchanged. Similarly, constitutive expression (overexpression) of a gene can be represented by setting the value of the associated node to *1*. For example, to investigate the overexpression of *HEY2*, we fix its value to *1* (i.e. *x*_*HEY2*_ (*t* + *1*) = *1*). We refer to the BN where the underlying perturbation is fixed as *perturbed BN*. To determine whether the knockout/overexpression is successful or not, we check how the knockout/overexpression alters the steady state landscape of the BN. In other words, we compare the steady state configuration of the original BN with that of the perturbed BN.

#### Results

We present the perturbations in Table 4. The first column is the *identifier (id)* of the perturbation while the second column contains the *perturbed node(s)* which can be either a knockout or an overexpression. The third column includes the *literature* where the corresponding perturbation is referenced while the fourth column declares whether the BN model successfully reproduced the knockout.

**Table 4:** The model replicates 5 out of 7 perturbations found in the literature.

| *id* | *Perturbed Node(s)* | *Literature* | *Success* |
| --- | --- | --- | --- |
| (i) | *HAND2* | *HAND2* knockout results in no right ventricle [42, 36, 35]. | Yes. |
| (ii) | *NR2F2* | Gene *NR2F2* encodes *COUPTF-II* transcription factor. *COUPTF-II* knockout results in no atrial cardiomyocytes [27]. | Yes. |
| (iii) | *TBX5* | *TBX5* knockout results in no atrial cardiomyocytes [6]. | No. |
| (iv) | *NKX2.5* | *NKX2.5* knockout results in no left ventricle [3]. | Yes. |
| (v) | *NKX2.5, HAND2* | *NKX2.5* with *HAND2* knockout gives no ventricle [42]. | Yes. |
| (vi) | *HEY2* | *HEY2* knockout mouse cannot develop a normal left ventricle. *HEY2* knockout mice get ectopic atrial gene expression in the left ventricle [26, 15]. | No. |
| (vii) | *HEY2* | Overexpression of *HEY2* in atrial cardiomyocytes represses atrial genes [26, 14, 41]. | Yes. |

In perturbation *(i)*, the knockout of *HAND2* results in a perturbed BN that exhibits six steady states (just like the original BN), but in all of them the ventricular marker *MYL2* is *off (0)* which means that we do not get ventricular cardiomyocytes at all. Therefore, we consider that our model successfully reproduces this knockout. In the case perturbation *(ii)*, the knockout of *NR2F2* results in a perturbed BN that exhibits again six steady states. In all of them, the atrial marker *MYL7* is *off* and, therefore, we consider this perturbation successful. The knockout of *TBX5* (perturbation *(iii)*) results in six steady states which are exactly the same as in the original BN but, in all of them, the node *TBX5* is *off*. This leads us to consider that the perturbation is unsuccessful; there exist steady states where the atrial marker *MYL7* is still expressed. In other words, the perturbed node (*TBX5*) does not change the steady state configuration of the BN other than its own value in the steady state configuration. The knockout of *NKX2*.*5* (perturbation *(iv)*) results in a perturbed BN which exhibits six steady states. Our model successfully reproduces this knockout because the ventricular marker *MYL2* is *off* in all these six steady states. The same behaviour exhibits the perturbed BN in case *(v)*. The knockout of *HEY2* (perturbation *(vi)*) was not successful because the perturbed BN reproduces six steady states wherein the ventricular marker *MYL2* is still active in some of them. The overexpression of *HEY2* (perturbation *(vii)*) results in a BN with six steady states wherein the atrial marker *MYL7* is inactive in all of them.^3^

#### Interpretation

Overall, our model successfully reproduced *5* out of *7* perturbations. The knockout of *TBX5* (perturbation *(iii)*) is not successful because *TBX5* may not affect the atrial marker *MYL7* but other atrial markers like *ANF* and *CX40* [6]. Similarly, the knockout of *HEY2* (perturbation *(vi)*) is unsuccessful because *HEY2* may not repress the ventricular marker *MYL2* but other ventricular markers; the ones referenced in [26, 15]. Indeed, in *HEY2* knockout mice, atrial genes *ALC-1* and *MLC-2a* were expressed ectopically in left ventricle [15].

### 4.2 Probabilistic Simulations of the Merged Boolean Network

Due to the non-deterministic nature of our framework, we cannot assess the reachability of the merged BN towards the six biologically-relevant steady states which correspond to cardiomyocytes. To investigate the ability of our model in transitioning initial states to the biologically relevant steady states, we conduct simulations. We consider a probabilistic version of the BN [33] wherein, in order to transit from a predecessor state to the immediate successor state, only one variable of the BN is selected at random with equal probability for updating. For instance, consider the model with three variables of Example 2: the probability of each transition is *1* /*3* because each one of the *3* variables is uniformly selected for update with equal probability. It is then straightforward to see that the STG forms a Markovian Chain as shown in Fig. 7. For the case of the merged BN, which has *21* variables, the probability of each transition in the corresponding Markovian Chain equals to *1* /*21*.

**Fig. 7:**
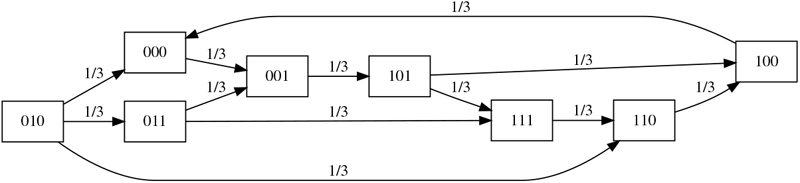
The Markovian Chain generated by uniformly selecting only one variable for update. Each state has exactly *3* outgoing transitions. The self-transitions of the states are not displayed in the figure for clarity. In contrast to the non-deterministic STG of Fig. 4 where the BN is in several states at a particular time point, here the BN is in exactly one state with some probability.

#### Experimental Design

We examine the behaviour of overall *400*.*000* independent and isolated cells: *100* .*000* for each of the *4* different activity combinations of the input variables (signals) *x*_*ex*_*WNT*_ and *x*_*RA*_ (see the first column of Table 5). The initial values of all the other variables is assigned with a random binary value in {*0, 1*}.

**Table 5:** Simulations.

| Inputs | FHF aCM | FHF left vCM | SHF aCM | SHF right vCM | null 1 | null 2 |
| --- | --- | --- | --- | --- | --- | --- |
| $x_{ex\_WNT} = 0, x_{RA} = 0$ | 0 | 81.937 | 0 | 0 | 18.163 | 0 |
| $x_{ex\_WNT} = 1, x_{RA} = 0$ | 0 | 0 | 0 | 100.000 | 0 | 0 |
| $x_{ex\_WNT} = 0, x_{RA} = 1$ | 81.858 | 0 | 0 | 0 | 0 | 18.142 |
| $x_{ex\_WNT} = 1, x_{RA} = 1$ | 0 | 0 | 100.000 | 0 | 0 | 0 |

#### Results

We present the results of the conducted simulations in Table 5. The first column (*Inputs*) contains the combination of the input variables (*x*_*ex*_*WNT*_, *x*_*RA*_) that each cell is treated with. In the first row (*x*_*ex*_*WNT*_ = *0, x*_*RA*_ = *0*), we investigate a population of *100* .*000* isolated and independent cells whose *WNT* and *RA* pathways are inactivated. Here, *81*.*937* out of *100*.*000* cells (more than *80%*) are differentiated into left ventricular cardiomyocytes of the first heart field, as expected. In the second row (*x*_*ex*_*WNT*_ = *1, x*_*RA*_ = *0*), we investigate the behaviour of the cells when *WNT* signalling pathway is activated while *RA* pathways remain inactive. In this case, all cells are differentiated into *SHF* right ventricular cardiomyocytes. The results of the *3rd* and the *4th* row of Table 5 can be interpreted accordingly.

#### Interpretation

We observe that *363*.*695* out of *400*.*000* simulated cells (*90*.*92%*) end up to the biologically relevant states. Importantly, the results are in agreement with in vitro protocols for cardiomyocyte differentiation [18]: for *x*_*ex WNT*=_ *0* (*1st* and *3rd* row of Table 5) we mostly obtain *FHF* cells which, when treated with RA (*x*_*RA*_ = *1*), give rise to *FHF aCMs*. On the contrary, for *x*_*ex*_*WNT*_ = *1* (*2nd* and *4th* row of Table 5) we always obtain a *SHF* cell which, in absence of *RA* signaling (*x*_*RA*_ = *0*), gives rise to an *SHF vCM*.

## 5 Discussion

Our study covers the entire lineage from an early cardiac progenitor towards chamber-specific cardiomyocytes. Initially, we present an interaction graph that describes cell differentiation of committed cardiomyocytes towards ventricular and atrial cardiomyocytes. The interaction graph has been coupled with Boolean dynamics whose correctness was validated in terms of its steady states; i.e. different steady states correspond to different cell types based on marker genes. After merging the cardiomyocyte BN with an existing BN that characterizes heart field formation, we identify steady states that correspond to cardiomyocytes located in specific cardiac regions. The investigation of the merged model is the first to describe the molecular mechanism for patterning heart-field-specific atrial and ventricular CMs. Particularly, the analysis of the merged model highlight the ability of the model to reproduce known knockout and overexpression experiments found in the literature, while the probabilistic simulations shows behaviours that are in agreement with in vitro differentiation protocols.

Extension of the merged BN with other dynamical formalisms, such as coupling it to the continuous domain, allows for further insights into the differentiation process. With this, it is possible to investigate the ratio-dependent dynamics of the signaling, as shown in Appendix C. We note that with the objective of keeping the model simple, several regulatory interactions are missing. Future work may focus on incorporating these missing interactions and adding other cell types and regulatory elements relevant to cardiac development. With this, the model would be able to reproduce all knockout experiments and allows for finding combinations of perturbations that may lead to more effective cardiomyocyte specification.

## Acknowledgements

Special thanks to the anonymous reviewers of the Conference *Computational Modeling in System Biology 2024* for their critical reviews. The study has also benefited from discussions with the following: Stefano Schivo from Open University of the Netherlands, Hil Meijer, Lucas F. Jansen Klomp, Xinqi Yan from the University of Twente, and Rebecca R. Snabel and Gert Jan C. Veenstra from Radboud University.

## A Developmental Origins of Cardiomyocytes

Fig. 8 provides a schematic overview of the developmental origins for CM commitment. Cardiac mesoderm is induced by expression of early cardiac transcription factor *MESP1* [28, 43]. Early cardiac progenitors, expressing *ISL1*, contribute to either the first heart field (FHF) or the second heart field (SHF). The heart fields can be distinguished by their characteristic marker expression. The FHF progenitors are identified by the lack of *ISL1* expression, in contrast to SHF progenitors which do express *ISL1*. The FHF generates the majority of left ventricular CMs, as well as part of the atrial. The SHF generates primarily right ventricular CMs, some part of atrial CMs and the outflow tract.

**Fig. 8:**
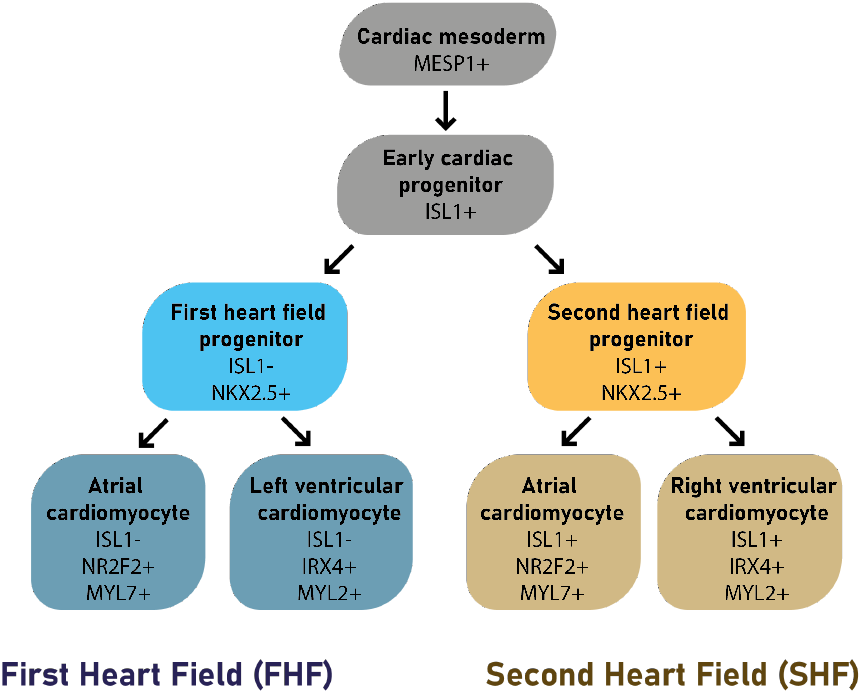
A more detailed representation of the differentiation process which includes marker genes of the specific cell states.

## B Supplementary Information of the Graph

**Fig. 9:**
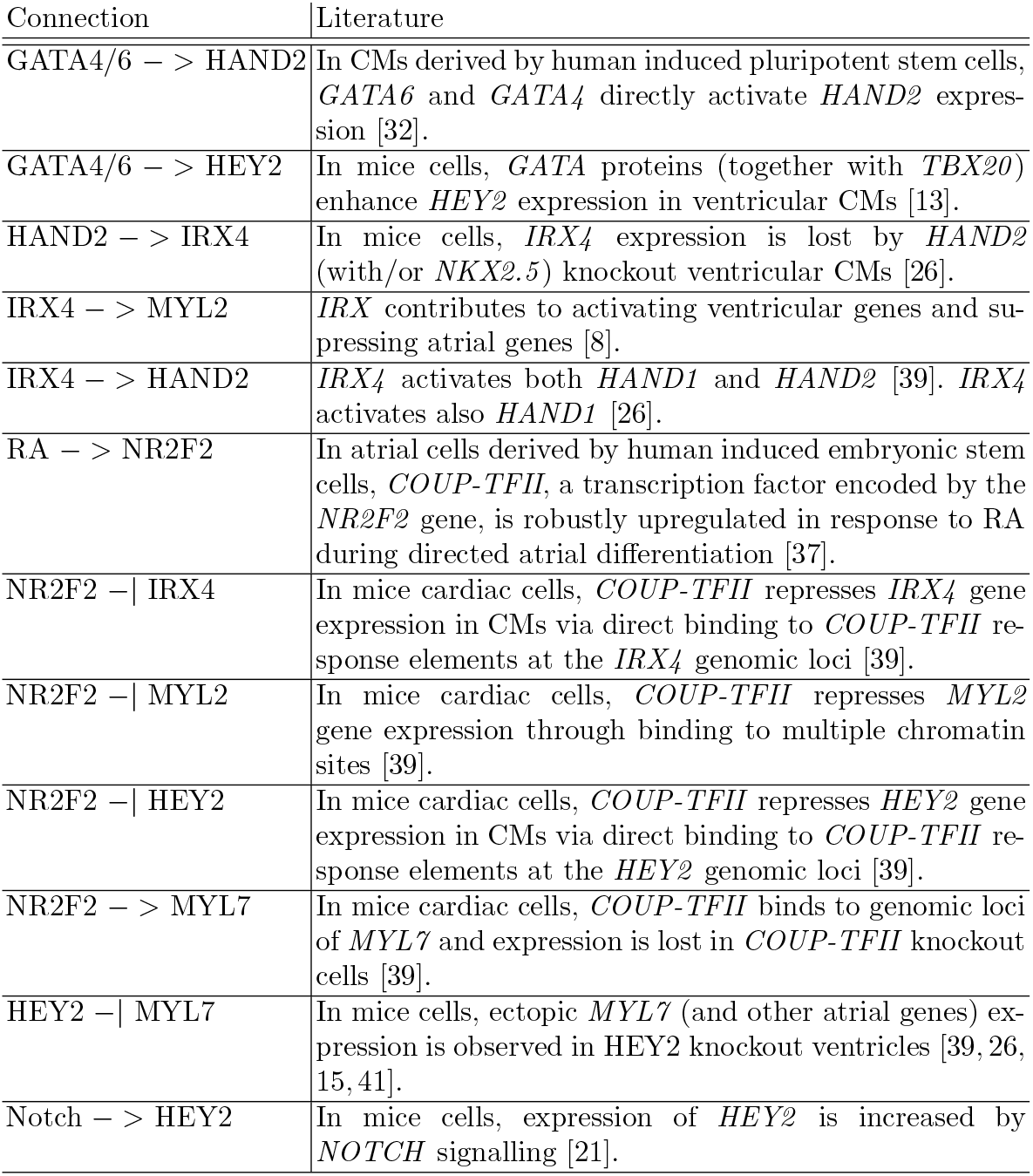
Extended evidence for the connections of the GRN in Fig. 2.

## C Transformation of the BN to other Formalisms

To further investigate the dynamical behaviour of the constructed BN, we reformulate to different dynamical formalisms [10, 16, 38]. Particularly, in Section C.1 we extend the domain of the BN variables to obtain another model called *Boolean Network Extension (BNE)* [10] where values lie in a continuous interval, in Section C.2 we convert the BNE to a continuous time analogue, and in Section C.3 we discuss related work.

### C.1 Extension to Real Dynamics and Analysis

To investigate the robustness of the model, we extend the BN to allow the variables to obtain values in the continuous interval [*0, 1*] as in [10] and then proceed to further analysis of the steady states of the model. The transformation of the Boolean rules into real-valued functions is the following:

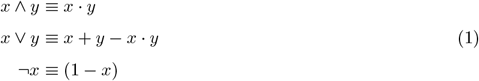

The left and the right hand side of the three equivalences of (1) provide the same results when *x, y* ∈ {*0, 1*}. Interestingly, the functions on the right hand side can mimic all the behaviors of the left hand side when the former is initialized with *0, 1*. On the other hand, when variables of the right hand side are initialized in [*0, 1*], they exhibit some more behaviors that the left hand side cannot reproduce. By applying the transformation rules to a BN, we obtain a discrete time dynamical system that we call Boolean Network Extension (BNE) as in [10]. We present the BNE of a simple BN in the following example:

#### Example 2

We consider the BN in the left part of Fig. 4 where x, y, z are the 3 Boolean variables with update functions x(t + 1) = y(t) ∨ z(t), y(t + 1) = x(t) ∧ z(t), and z(t + 1) = ¬x(t). The corresponding real discrete time dynamical system has the following form:

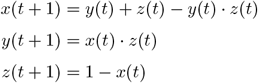

By applying the transformation rules to the BN of Fig. 2, we obtain the following BNE:

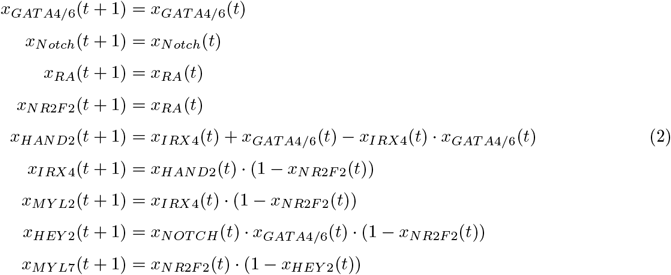

For the BNE, it is straightforward to see that if the variables are initialized with values in [*0, 1*] then the updated values will again lie in the same interval. The question that naturally arises is: does the BNE exhibit steady states other than the ones found in the original BN?

#### Parameter Definition and Steady States

Now, we proceed by finding the steady states of the variables. We set:

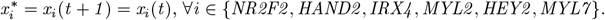

The BNE is the following:

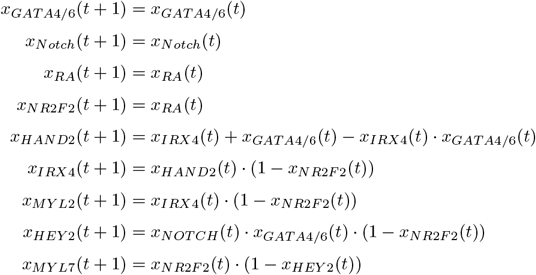

We propose that the input nodes *GATA4/6, RA, Notch* serve as parameters. We denote the parameter *x*_*GATA4/6*_ with γ, the parameter *x*_*RA*_ with ρ, and the parameter *x*_*Notch*_ with *ν*. We further justify the parameterization of these variables in the next section (ODE modelling of Section C.2).

Note that several variables are stabilized with respect to the parameters; the update function of *x*_*NR2F2*_ takes the form *x*_*NR2F2*_ (*t* + *1*) = *ρ*, and, consequently, the update function of *x*_*HEY2*_ takes the form *x*_*HEY2*_ (*t* + *1*) = *ν* · γ · (*1* − *ρ*). Indeed, the inputs *GATA4/6, Notch* and the stabilized *NR2F2* variable are the only incoming influences of the *x*_*HEY2*_ in the interaction graph. Furthermore, the stabilization of *x*_*HEY2*_ leads to consequent stabilization of *x*_*MYL7*_ with x_*MY*_*L*7_(t+1) = *ρ*·(1−(*ν* ·γ ·(1−*ρ*))). The variables that do not stabilize are {*x*_*HAND2*_, *x*_*IRX4*_, *x*_*MYL2*_} and their update functions are the following:

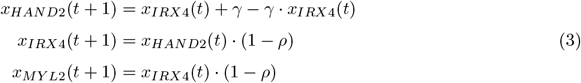

We proceed by finding the steady states of the variables. We set:

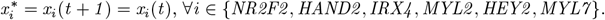

The stabilized variables remain on their fixed values by the parameters. For the variables of (3), we have:

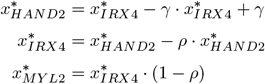

The latter is a linear system of equations. By solving the system, we obtain the variables’ steady states which depend only on the values of the parameters.

Overall, the steady states are the following:

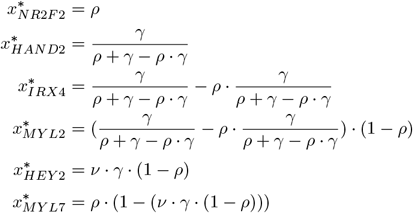

Interestingly, the steady states depend only on values of the parameters.

#### Steady State Analysis of the BNE

To investigate the effect of the parameter *ρ* on the steady states of the models, we fix the parameters γ = *ν* = *1* as long as the genes *GATA4/6* and *Notch* are expressed in the progenitors and remain expressed until all stages of heart development. In Fig. 10, we present the stabilized expression of the nodes {*NR2F2, MYL7, MYL2, IRX4, HEY2, HAND2*} when γ = *ν* = *1*, and *ρ* varies. We see that for high values of *RA* (> *0* .*5*) the steady state activity of *MYL7* is also increased and it is larger than the steady state activity of *MYL2* (i.e.,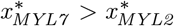), suggesting that the cell is an atrial CM (see blue and yellow line of Fig. 10). In contrast, for low values of *RA* (< *0* .*5*), the steady state activity of *MYL2* is higher than that of *MYL7* (i.e., 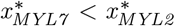), suggesting that the cell is a ventricular CM.

**Fig. 10:**
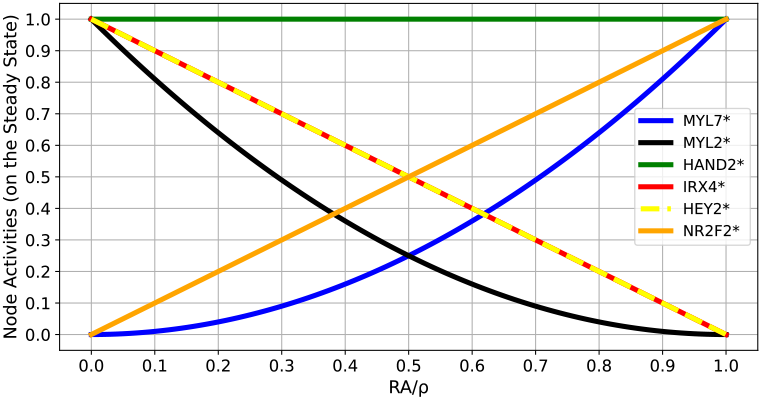
Steady state node activities upon RA administration. The X-axis is the value of 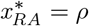. The Y-axis is the activity of nodes on their steady states.

While the value of *RA* is increasing, the steady state activities of *HEY2* and *IRX4* are decreasing. On the contrary, the steady state activity of *NR2F2* increases linearly. The latter is reasonable considering that *HEY2, IRX4* are ventricular specific markers while *NR2F2* is atrial specific. However, the node *HAND2* remains fully active on all steady states, independent of the value of *RA*. The latter is a consequence of the parameter γ (i.e. the activity of *GATA4/6*) which remains stable and equal to *1*.

### C.2 Transformation to Continuous-time ODE Model

Transformation from BNs to continuous time ordinary differential equations [16, 38] do exist. The transformation that we perform in this section can be considered to be the continuous time analogue of the BNE. Next, we provide the transformation of the basic Example 2:

*Example 3*. Considering the BN of with update functions x(t + 1) = y(t) ∨ z(t), y(t + 1) = x(t) ∧ z(t), and z(t + 1) = *¬*x(t), the corresponding ODE system has the following form:

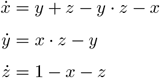

One can easily see that the ODE system is the abstraction of the BNE with respect to time (see Example 2). For the BN of Fig. 3, the full set of ODEs is the following:

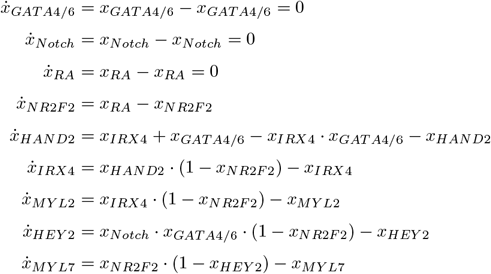

Note that the time evolution 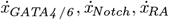 of these variables equals zero which motivates us to consider these variables as parameters, like we did in Section C.1.

### C.3 Related work

Our approach for dynamical modelling of Section 2.2 constitutes a generic modeling pipeline. Several parts of the dynamical modelling can be reproduced by existing software. In Section 2.2, we constructed a BN model with respect to a prescribed steady state configuration and an interaction graph (first box of Fig. 11). Although the cardiomyocyte BN was developed by hand, BN tools like Sketches [2] and Griffin [22] (2nd dashed box of Fig. 11) can automatically generate a BN (3rd box of Fig. 11) based on a given interaction graph and a prescribed attractor configuration. Later, we utilize the Boolean Network Extension [10] to further analyze our model (upper right box of Fig. 11). According to [19], this is a probabilistic generalization of Boolean logic that George Boole himself had proposed. The BNE is the simplest discrete time analogue of the Odefied BN [16]. Nevertheless, the transformation by Odefy (box *Ordinary Differential Equations* on the bottom right part of Fig. 11) requires and provides the ability for further parameterization. The extra parameters may characterize, for example, the timescales of the updates, or the required level of activation when the update functions of the nodes have the form of Hill functions. Molecular interactions are known to show a switch-like behavior, which can be modeled using sigmoid-shaped Hill functions and whose parameters determine the sigmoid shape [12, 38]. SQUAD software [20] also transforms BNs to continuous-time and continuous-space systems.

**Fig. 11:**
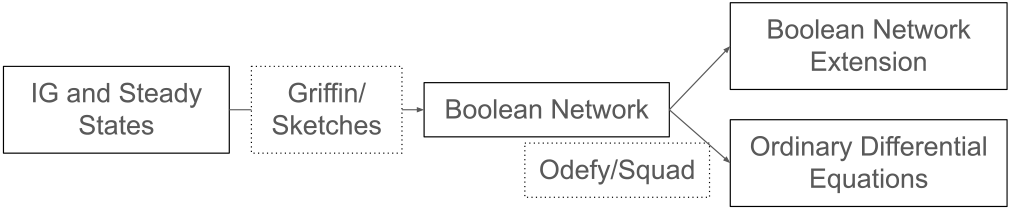
A toolchain for performing the analysis of Section 2.2.

## D The Merged Boolean Network

The merged BN consists of the heart field BN and the cardiomyocyte BN. Changes are marked with **bold**.

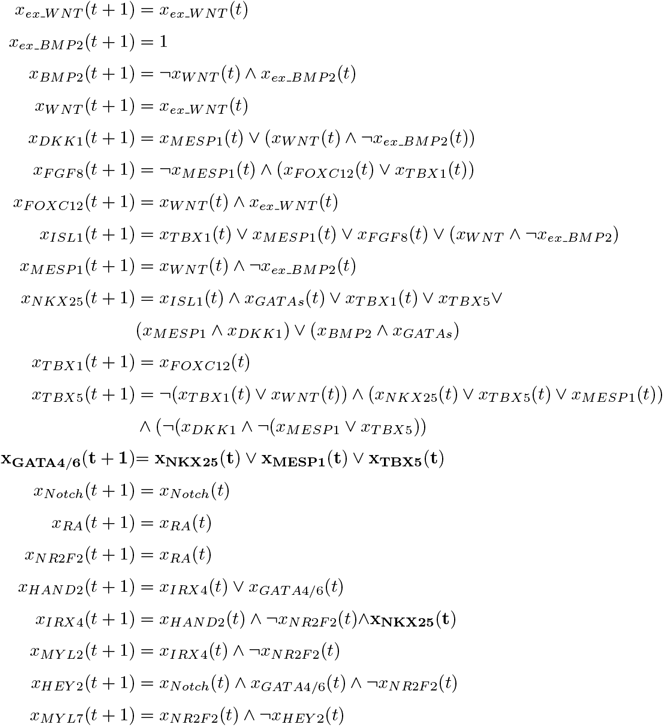

## Footnotes

1 The results of our study are reproducible by CoLoMoTo [24] notebooks: github.com/GeorgiosArg/Beyond-Committed-CMs.

2 We perform further analysis of this BN in Appendix C where we transform it to other dynamical formalisms.

3 The results of these and other perturbations can be found on: github.com/GeorgiosArg/Beyond-Committed-Cardiomyocytes.

